# Learning to see the Ebbinghaus illusion in the periphery reveals a top-down stabilization of size perception across the visual field

**DOI:** 10.1101/2020.04.05.026062

**Authors:** Cécile Eymond, Tal Seidel Malkinson, Lionel Naccache

## Abstract

Our conscious visual perception relies on predictive signals, notably in the periphery where sensory uncertainty is high. We investigated how such signals could support perceptual stability of objects’ size across the visual field. When attended carefully, the same object appears slightly smaller in the periphery compared to the fovea. Could this perceptual difference be encoded as a strong prior to predict the peripheral perceived size relative to the fovea? Recent studies emphasized the role of foveal information in defining peripheral size percepts. However, they could not disentangle bottom-up from top-down mechanisms. Here, we revealed a pure top-down contribution to the perceptual size difference between periphery and fovea. First, we discovered a novel Ebbinghaus illusion effect, inducing a typical reduction of foveal perceived size, but a reversed increase effect in the periphery. The size percept was similar at both retinal locations and deviated from the classic perceptual difference. Then through an updating process of successive peripheral-foveal viewing, the unusual peripheral perceived size decreased. The classic perceptual eccentricity difference was restored and the peripheral illusion effect changed into a fovea-like reduction. Therefore, we report the existence of a prior that actively shapes peripheral size perception and stabilizes it relative to the fovea.

## Introduction

Perception is a complex process in which active mechanisms infer from sensory signals the content of our visual awareness. One of the most intriguing questions is how such mechanisms ensure the impression of a uniformly rich visual scene, while sensory information becomes drastically coarser with eccentricity ^1,2^. Several mechanisms have been found to resolve sensory uncertainty in the periphery by combining the poor incoming information with predictions based on our highly detailed visual experience at the fovea ^3–7^. Still our conscious perception in the periphery remains limited compared to central viewing and we can easily notice it by attending carefully to visual features of peripheral objects.

In the present study, we investigated another potential role of predictive mechanisms that shape our conscious peripheral perception. We focused on object size, which appears slightly smaller in the periphery compared to the fovea ^8–10^. What can such size underestimation tell us about predictive mechanisms subtending peripheral perception? In addition to coping with the heterogeneity of sensory input, how do predictive mechanisms support perceptual stability across the visual field? We tested the existence of a top-down contribution to the perceptual size difference between periphery and fovea. Such a mechanism would support a stable perception of size across the visual field, which is crucial to guide efficiently our interactions with the surrounding world.

We report two results. We first discovered a novel illusory effect in the visual periphery. When the Ebbinghaus illusion ^11,12^ is presented at the fovea, the same disk is perceived larger or smaller depending on the contextual surrounding disks – the inducers ^13,14^. We found specific inducers eliciting a reduction of perceived size in the fovea, and a reversed increase effect in the periphery. This Ebbinghaus disk thus deviated from the classic perceptual bias of a decrease in perceived size with eccentricity and was perceived as large in the periphery as in the fovea. We hypothesized that if the difference in perceived size between periphery and fovea is a key aspect of perceptual stability, the deviating peripheral percept would be updated to better match this difference.

Second, we confirmed the existence of a top-down contribution to the perceptual size difference between periphery and fovea. We used a recent method showing how the peripheral size percept is adjusted relatively to foveal viewing ^7,15^. Valsecchi and Gegenfurtner showed that the same disk was perceived as smaller or larger in the periphery after being associated with a smaller or larger disk presented in the fovea. In each trial, participants first reported the perceived size of a peripheral disk and then viewed it centrally, for instance by making a saccade towards it. When the peripheral disk was replaced during the saccade by a slightly different disk that was viewed consequently in the fovea, the perceived size of the peripheral disk changed accordingly through successive trials. Without replacement during the saccade, the perceived size of the peripheral disk remained unchanged, and the authors observed the classic perceptual size difference, with the disk perceived smaller in periphery relative to fovea.

In these previous studies ^7,15^, the size of the stimulus presented in the fovea differed physically from the one in the periphery. This sensory difference in the fovea may have driven the change in peripheral perception through a bottom-up effect. We adapted this peripheral-foveal contingency paradigm and tested its effect on the peripheral perceived size of our Ebbinghaus disk, which kept a constant physical size at both retinal locations. We observed a decrease in peripheral perceived size, suggesting that the peripheral percept was updated to better match the classic perceptual difference with fovea. This result demonstrates the existence of a pure top-down contribution to the peripheral-foveal difference that may underlie perceptual stability across the visual field.

## Results

### Experiment 1

In this experiment we showed that the Ebbinghaus illusion can be used to bias the classic perceptual size difference between periphery and fovea. We found specific inducers that altered the perceived size of a black disk differently in the periphery and in the fovea. We used a comparison task to evaluate how large the participants perceived our Ebbinghaus disk at both retinal locations and compared the results to those of an isolated disk (Fig. 1a-c). We repeated the procedure for three Standard disks (1.5°, or 1.6°, or 1.7°-diameter, see Methods) and analyzed the main and interaction effects on the apparent size, defined as the physical size of a Test disk that should be displayed to be perceived as large as the Standard disk (as indexed by the point of subjective equality (PSE), see Methods). A decrease in apparent size between the two retinal locations, or disk types, would reflect a smaller perceived size.

**Figure 1.**
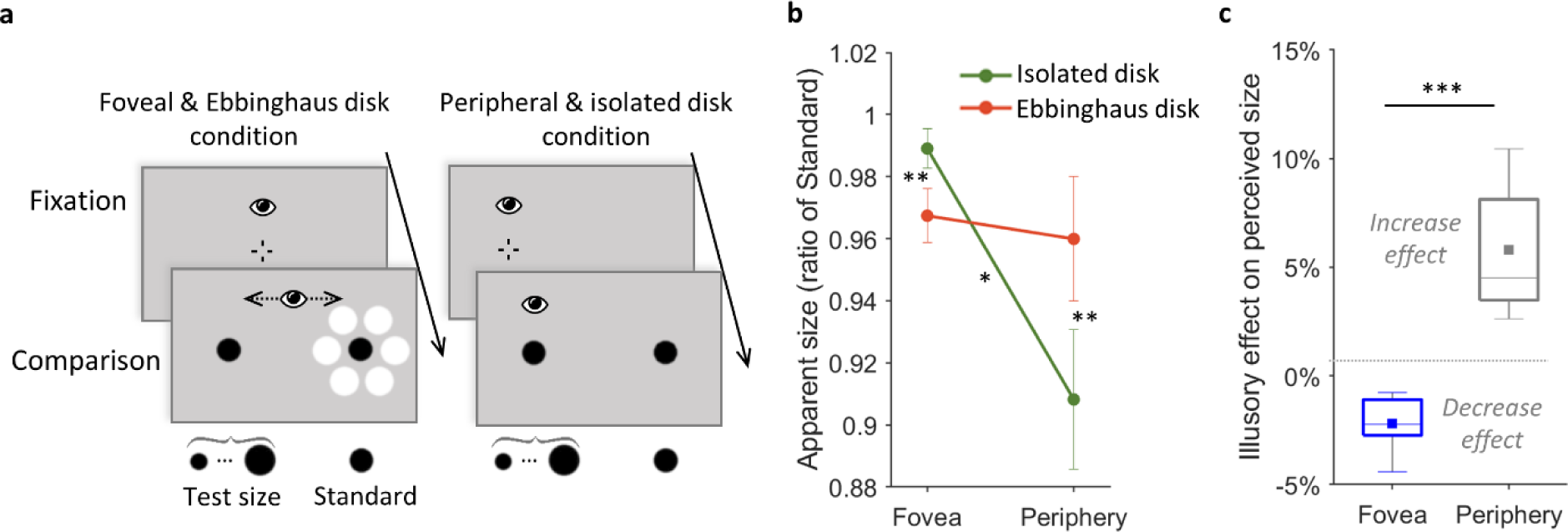
Experiment 1. (a) Experimental procedure. The Standard black disk was always displayed on the right side of the screen, isolated (right panels) or as an Ebbinghaus disk (left panels). The perceived size of each disk type was evaluated at the fovea (left panels) and in the periphery (right panels). Each trial started with a fixation at the screen center (illustrated on the left side in the peripheral condition for convenience). A Test disk was presented on the screen left side in the foveal condition, or replaced the fixation cross in the peripheral condition. Participants had to answer which of the two black disks they perceived the largest. In the foveal condition, participants could freely look overtly at both stimuli during the comparison task. In the peripheral condition, participants maintained the central fixation all along the trial. In these illustrated trials, all black disks have the same physical size. Stimuli are not drawn to scale. (b) Results: Mean apparent size for the isolated disk (green line) and the Ebbinghaus disk (red line), viewed foveally or at 12° to the right periphery. The apparent size of the Standard disk was defined as the physical size that the Test size should have to be perceived as large as the Standard disk. It was evaluated as a ratio of the Standard disk size and averaged across the three tested Standard disk sizes (see Table 1 in Methods section). A smaller (larger) apparent size indicated a smaller (larger) perceived size. Error bars indicate ± SEM (c) Results: Boxplots of the illusory effect on perceived size, defined as the percentage of apparent size difference between the Ebbinghaus and isolated disks, in the foveal (blue boxplot) and in the peripheral conditions (grey boxplot). Squares represent mean illusory effects. Negative values indicate a decrease effect; positive values an increase effect on perceived size. (B)-(C) * p<.05, ** p<.01 and *** p<.001

A 2⨯2⨯3 repeated measures ANOVA revealed a significant interaction between the disk type (Ebbinghaus and isolated disks) and the retinal location (peripheral and foveal) factors (see Fig. 1b). [F(1,6)= 68.91, p=.0002, η_p_^2^= .920]. Before analyzing further this interaction, we confirmed that the main effect of the Standard size (1.5°, or 1.6°, or 1.7°-diameter) was not significant [F(2,12)= 1.60, p=.244], as well as its interactions with the other factors [Standard size x retinal location: F(2,12)= 0.98, p=.403; Standard size x disk type: F(2,12)= 0.16, p=.857; and Standard size x retinal location x disk type: F(2,12)= 1.20, p=.334]. The Bayesian analysis confirmed that the best model fitting the data compared to the null model (BF_10_= 5.10^6^, error 6.11%) included the main effect of retinal location (periphery or fovea, BF_incl_= 4708.852), the main effect of disk type factor (isolated or Ebbinghaus disk, BF_incl_=0.796), and their interaction (BF_incl_= 1704.617). The Standard size (BF_incl_=0.198), as well as its interaction with others factors, had lower inclusion Bayesian factors in the analysis of effects (Standard size x retinal location BF_incl_=0.187; Standard size x disk type BF_incl_=0.167; Standard size x retinal location x disk type BF_incl_=0.326). Therefore, for further analyses, the mean apparent size for each disk type and retinal location was computed across Standard sizes.

#### Reversed illusory effect in the periphery

The mean illusory effect at the fovea was a -2.2% reduction of perceived size (95% CI [-3.3, - 1.0], one-sample two-tailed t(6)=-4.63, p=.004, d=-1.750; Fig. 1c), replicating the typical decrease induced by such contextual modulation in central vision ^13,14^. However, in the periphery the inducers had a reversed effect (pair-wise two-tailed t(6)=7.85, p=.0002, d=2.966), eliciting a 5.8% mean increase of perceived size (95% CI [3.1, 8.5], one-sample two-tailed t(6)=5.24, p=.002, d=1.982; Fig. 1c).

#### Lack of perceptual size difference between periphery and fovea

Overall, the perceived size of the isolated disk did match the classic decrease with eccentricity ^8–10^, with a mean peripheral-foveal apparent size ratio of 0.92 significantly smaller than 1 (95% CI [0.86, 0.98], one-sample two-tailed t(6)=3.34, p=.016, d=1.262; Fig. 1b]). However, the disk surrounded by inducers was perceived as large in the periphery as in the fovea, with an apparent size ratio of 0.99 (95% CI [0.94, 1.05], two-tailed one-sample against 1, t(6)=0.32, p=.759; BF_01_=2.711, error 4.10^−5^ %; Fig. 1b) significantly different from that of the isolated disk (pair-wise two-tailed t(6)=8.47, p=.0002, d=3.201; Fig. 1b). This revealed for our Ebbinghaus disk an unusual deviation from the classic peripheral-foveal perceptual size ratio.

To sum up, the results of Experiment 1 replicated for the isolated disk the classic decrease in perceived size with eccentricity. We observed the Ebbinghaus illusion effect in the fovea, where the inducers reduced the perceived size of the black disk. Notably, the inducers had a reversed effect in the periphery and increased the perceived size of the black disk. The size of the Ebbinghaus disk was thus perceived similarly at both retinal locations. We hypothesized that if the classic perceived size difference between periphery and fovea is a key aspect of perceptual stability, the peripheral perceived size of the Ebbinghaus disk would be updated to better match this difference, whereas that of the isolated disk would remain stable.

### Experiment 2

In Experiment 2 we tested our hypothesis about a top-down contribution to the perceptual size difference between periphery and fovea. We examined whether the perceived size of our Ebbinghaus disk in the periphery, which initially did not differ from its foveal view (Experiment 1), would be updated to solve this lack of perceptual difference. Notably, in Experiment 1 we evaluated peripheral and foveal perception independently (see Methods). In Experiment 2, we introduced a peripheral-foveal contingency paradigm, adapted from a procedure in which changing an object’s physical size only when viewed foveally changes accordingly its peripheral perceived size ^7^. In each trial, while keeping constant the stimulus physical size, we first asked the participants to perform a comparison task to evaluate how large they perceived the stimulus in the periphery, then asked them to make a saccade towards it, establishing so the peripheral-foveal contingency (Fig. 2a). They repeated this trial procedure nearly 300 times with the Ebbinghaus disk in one session, and with the isolated disk in another session. We compared the peripheral perceived size of both disk types between the pre- and post-test phases, comprising respectively the first and last hundred trials (Fig. 2a & 2b). The perceived size was evaluated with the equivalent size, defined as the physical size that the peripheral Test disk must have to be perceived as large as a foveal Reference (as indexed by the PSE see Methods). An increase in the peripheral equivalent size between the pre- and post-tests indicated a decrease in the peripheral perceived size.

**Figure 2.**
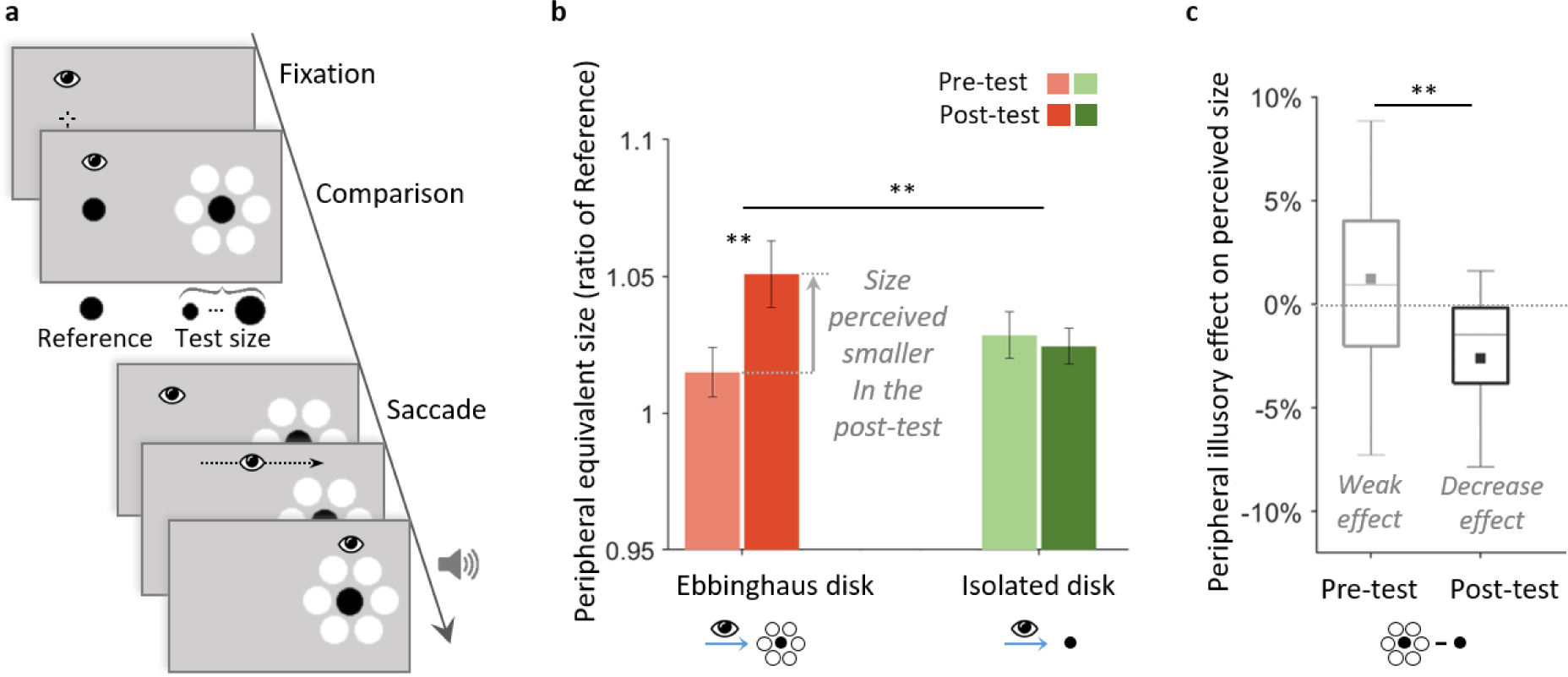
Experiment 2, main group, with peripheral-foveal contingencies. (a) Trial procedure. While maintaining central fixation, participants first compared the foveal Reference disk to a peripheral black Test disk that was either isolated or an Ebbinghaus disk (as illustrated), in two separate sessions. After their response, the foveal Reference disappearance signaled the participants to make a saccade towards the peripheral Test disk. Participants received an auditory feedback on landing accuracy: a high tone indicated an accurate saccade (see Methods section for criterion detail) and a low tone that the accuracy might be improved. During a trial, stimuli kept a constant physical size. All trials followed this procedure, the first and last third of the total of trials constituted the pre- and post-test phases. Stimuli are not drawn to scale. (b) Results: Mean peripheral equivalent size of the foveal Reference in the pre- and post-tests, for the isolated (green bars) and the Ebbinghaus disks (red bars) conditions. The peripheral equivalent size was defined as the physical size that the peripheral Test disk must have to be perceived as large as the foveal Reference. It was evaluated as a ratio of the foveal Reference (see Methods). An increase in the peripheral equivalent size between the pre- and post-tests indicated a decrease in the peripheral perceived size: the Test disk had to be larger in the post-test to match the perceived size of the foveal Reference. Error bars indicate ± SEM. (c) Results: Box plots of the peripheral illusory effect in the pre- and post-tests, defined as the percentage of perceived size difference between the Ebbinghaus and isolated disks conditions. Filled squared indicate the means. Negative values indicate a decrease effect, positive values an increase effect on perceived size. (b)-(c) ** p<.01

A 2⨯2⨯2 mixed ANOVA showed that the disk type (isolated vs. Ebbinghaus disk) and the evaluation phase (pre- vs. post-test) within-subject factors interacted significantly [F(1,22)=7.68, p=.011, η_p_^2^=.259; BF_incl_=5.443]. Before further analyzing this interaction, we confirmed that the disk type order had no significant main effect (the between-subjects factor: Ebbinghaus disk tested before vs. after the isolated disk; [F(1,22)=1.04, p=.319; BF_incl_=0.550]), as well as no significant interaction with the other factors [order x disk type: F(1,22)=2.68, p=.116, BF_incl_=0.778; order x phase: F(1,22)=2.24, p=.149, BF_incl_=0.632; order x disk type x phase: F(1,22)=1.44, p=.242, BF_incl_=0.924]. In all the following analyses, data were collapsed across disk type order.

#### Restored perceptual size difference between periphery and fovea

As hypothesized, the peripheral perceived size of the Ebbinghaus disk decreased by -3.6% between the pre- and post-tests phases (95% CI [-5.8, -1.4], one-sample two-tailed t(23)=3.39, p=.003, d=0.692; Fig. 2b), as indicated by the increase in the peripheral equivalent size. The peripheral Ebbinghaus disk had to be physically larger in the post-test compared to the pre-test to match the foveal Reference (see Methods). This decrease in perceived size between the pre- and post-test phases was larger than for the isolated disk (pair-wise two-tailed t(23)=2.84, p=.009, d=0.580), which perceived size did not change significantly (mean 0.3%, 95% CI [-1.0, 1.7], one-sample two-tailed t(23)=-0.48, p=.636; BF_01_=4.194, error 0.03%; Fig. 2b).

#### Updated illusory effect in the periphery

The perceived size of both disk types were not significantly different in the pre-test phase (mean difference 1.2%, 95% CI [-0.6, 3.1], one-sample two-tailed t(23)=-1.38, p=.181; BF_01_ =2.017, error 1.10^−4^ %). We compared this non-significant peripheral illusory effect to that in the post-test, where the presence of the inducers had an effect significantly larger (pair-wise two-tailed t(23)=2.79, p=.010, d=0.569; Fig. 2c). We found that the Ebbinghaus disk was perceived significantly smaller than the isolated disk in the post-test phase (mean -2.1%, 95% CI [-4.8, -0.4], one-sample two-tailed t(23)=2.48, p=.021, d=0.507; Fig. 2b). Therefore, the peripheral illusory effect changed into a reduction on perceived size, as in the fovea.

#### Control group without peripheral-foveal contingency

Finally, a 2×2 mixed ANOVA revealed that the updating for the Ebbinghaus disk observed in the main group with peripheral-foveal contingencies was different from that observed in a control group without peripheral-foveal contingency (see Fig. 3a & b). The effect of saccade execution (between-subjects factor) on peripheral equivalent size was significant [F(1,34)=7.18, p=.011, η_p_^2^=.174; BF_incl_=6.277] as well as the effect of the evaluation phase (pre-vs. post-test, within-subject factor) [F(1,34)=7.18, p=.011, η_p_^2^=0.174; BF_incl_=20.985]. The interaction between these two factors was not significant [F(1,34) =2.87, p=.099; BF_incl_=3.585]. Overall in the control group, the percentage of change in peripheral equivalent size between the pre- and post-test phases was not significant [mean 0.9%, 95% CI [-0.9, 2.6], one-sample two-tailed t(11)=1.05, p=.315; BF_01_=2.195, error 0.02%], suggesting that the peripheral perceived size of the Ebbinghaus disk was not updated without peripheral-foveal contingency.

**Figure 3.**
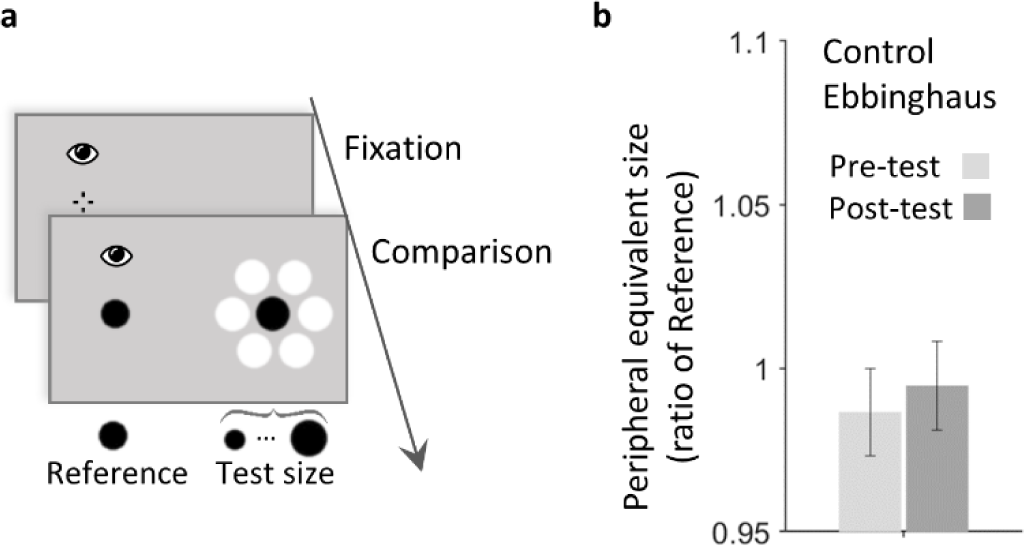
Experiment 2, control group, Ebbinghaus without contingency. (a) Trial procedure. While fixating, participants compared the foveal Reference disk to a peripheral black Ebbinghaus Test disk. All trials followed this procedure, the first and last third of the total of trials constituted the pre- and post-test phases. Stimuli are not drawn to scale. (b) Results: Mean peripheral equivalent size of the foveal Reference in the pre- and post-tests for the Ebbinghaus disk control condition. The peripheral perceived size was evaluated using the peripheral equivalent size, which was defined as the physical size that the peripheral Test disk must have to be perceived as large as the foveal Reference. It was evaluated as a ratio of the foveal Reference (see Methods). Error bars indicate ± SEM.

We also compared the perceived size of this control - Ebbinghaus disk group (Fig. 3b) to the results of the main - isolated disk group (Fig.2b, green bars). A 2⨯2 mixed ANOVA revealed that the main effect of the phase (pre- & post-tests) was not significant [F(1,34)=0.12, p=.727; BF_incl_=0.216], like its interaction with the factor group [F(1,34)=1.12, p=.298; BF_incl_=0.334]. However, the main effect of the group was significant [F(1,34)=7.53, p=.010, η_p_^2^=.181; BF_incl_=3.910]. Thus, the peripheral perceived size was larger in the control Ebbinghaus disk group than in the isolated disk group (both in the pre- and post-test phases), replicating the reversed illusory effect in the periphery observed in Experiment 1.

To sum up, in Experiment 2 we observed for the Ebbinghaus disk a decrease between the pre- and post-test phases in peripheral perceived size, which was initially similar to the one in the fovea (see Experiment 1). The perceptual difference between periphery and fovea was restored, and the peripheral illusory effect changed from the increase observed in Experiment 1 to a reduction on perceived size. The peripheral perceived size of the isolated disk that matched the classic perceptual difference with the fovea remained unchanged, similarly to that of the Ebbinghaus disk when not successively viewed in the periphery and the fovea.

## Discussion

In this study, we tested the existence of a top-down effect contributing to the perceptual size difference between periphery and fovea, which may support stability of size perception across the visual field. First, we report a novel Ebbinghaus illusion effect. Our Ebbinghaus disk was perceived as larger than an isolated disk when presented in the periphery, but smaller in the fovea. The peripheral illusion had thus an increase effect opposite to the typical reduction observed in the fovea, and the Ebbinghaus disk size was perceived similarly at both retinal locations. This first result showed therefore for our Ebbinghaus disk an unusual deviation from the peripheral-foveal perceptual size difference, while the isolated disk size perception matched the classic decrease with eccentricity.

We then tested the updating of this unusual peripheral percept through successive peripheral-foveal presentations of the Ebbinghaus disk. We observed a decrease of the peripheral perceived size, which suggests that the percept was updated to better match the classic perceptual difference with the fovea. The illusory effect in the periphery changed consequently into a reduction, like in the fovea. This second finding therefore reveals a pure top-down effect maintaining the peripheral percept smaller than the one in the fovea.

In Experiment 1 the peripheral illusion may had initially an unusual effect because our ability to recognize an object in a clutter is strongly affected in the periphery. The inducers would have elicited this crowding phenomenon^16^ which might have altered the peripheral illusory effect. The two main alternative mechanisms suggested for the foveal illusion ^17,18^ might both have interacted with crowding in the periphery. At low-level processing, interactions of adjacent contours have been suggested to underlie the foveal illusion^18^ or to contribute to the crowding phenomenon in the periphery^19^. Similarly, at higher stages of visual processing, crowding in the periphery was shown to harmonize features of the target to that of the flankers^20^ and being surrounded by larger inducers may have caused the peripheral black disk to seem larger - an assimilation process also suggested to explain the foveal illusion^21^. Additional factors such as luminance could explain the initial illusory effect observed in the periphery. The white inducers might have increased the luminance difference between the target disk and the background, which could have increased the perceived size in the periphery ^22^. The unusual illusory increase effect in the periphery may have counterbalanced the eccentricity dependent decrease in perceived size, leading to similar perceived size at both retinal locations. The results of Experiment 2 then showed that the typical reduction effect of the illusion was restored in the periphery through successive peripheral-foveal viewing of the Ebbinghaus disk.

In Experiment 2 for the Ebbinghaus disk, we expected a reduction in perceived size greater than the -3.6% mean decrease we observed between the pre- and post-test phases. We expected a mean decrease closer to -5.6%, an estimate based on Experiment 1 results suggesting that a -8% decrease would restore the classic perceptual size difference with fovea, and on the results of Valsecchi and Gegenfurtner (2016) who observed 70% of the expected change. This lack of updating was likely the outcome of the experimental procedure. Since a peripheral-foveal contingency took place in each trial, the updating already partially built up in the pre-test phase, leading to some decrease in the peripheral perceived size of the Ebbinghaus disk. This also explains why we did not observe in the pre-test phase a reversed illusory effect like in Experiment 1, but instead both the Ebbinghaus and isolated disk sizes were perceived similarly. Finally, we observed in the post-test phase of Experiment 2 an updated illusory effect in the periphery with a -2.2% reduction on perceived size that closely matched the illusory effect observed in the fovea in Experiment 1 with -2.1% reduction on perceived size.

To discard the alternative explanation of pure perceptual learning, we tested in a control condition whether the updating of the peripheral percept resulted from the successive viewing at both retinal locations. Indeed, perceptual learning could release crowding ^23,24^ and may have occurred through the repetition of the comparison task over successive trials. Such perceptual learning would have supported the emergence of the inducers’ reduction effect in the periphery. As expected, without the subsequent foveal view, the peripheral perceived size of the Ebbinghaus disk was not updated. Moreover, we replicated the peripheral reversed illusory effect observed in Experiment 1, when comparing the perceived size of this Ebbinghaus disk in the control condition to that of the isolated disk that was not updated between the pre- and post-test phases. This confirmed the critical role of successive peripheral-foveal viewing of the stimulus in the updating process.

Furthermore, the updating was specific to the Ebbinghaus disk which percept initially deviated from the classic peripheral-foveal size difference. Testing first the Ebbinghaus disk did not cause an updating in the following isolated disk condition. This discarded any learning of a general correction to the peripheral perceived size that would be further transferred to other stimuli, even to those matching the classic decrease in perceived size with eccentricity, regardless of the foveal input.

We did not control for the inducers’ size perception in the periphery, which might have changed between the pre- and post-test phases. This could have influenced the peripheral illusory effect in the comparison task through modulation of the size contrast^25^. We thus cannot discard this potential alternative explanation for the observed updating, even if very unlikely. We asked explicitly the participants to pay attention only to the black disk and not to the inducers in the comparison task, and similarly at saccade landing, we encouraged the participants to focus on the black disk using an auditory feedback on landing accuracy (checked within a 1°-circular region around the inner black disk). Additionally, according to the spontaneous subjective reports of some participants, they failed to perceive clearly the peripheral Ebbinghaus inducers even after the updating.

Taken together, our results support the key role of the peripheral-foveal perceptual difference in size perception stability. They show that predictive signals contribute to the peripheral percept by maintaining it smaller than in the fovea. This top-down contribution to the decrease in perceived size with eccentricity complements the contribution of low-level processing in early visual cortex shown recently ^10^. A high-level mechanism based on a strong prior according to which objects appear smaller in the periphery compared to the fovea may support perceptual size stability across the visual field. The discrepancy between the initial peripheral percept and the predicted percept formed by combining the foveal input and the smaller size prior may have driven the updating. This explanation is consistent with previous results^7^, when an updating of the peripheral size percept was observed after a foveal physical change: the predicted peripheral percept based on such a prior would have changed according to the foveal perceived size modulation.

The predictive aspect of peripheral perception was previously suggested to result from learned associations between the peripheral and foveal images of the same object ^3^. After learning a new association between two slightly different peripheral and foveal objects, the observed peripheral percept better matched the recently associated foveal input. Such associative learning supports the updating of an internal sensory model, which is at odds with a mechanism based on a strong prior. According to this associative learning account ^3^, we should not have observed any change in peripheral perception in order to converge with the foveal input, since the initial percept in our experiment was similar at both retinal locations. We can thus discard the exclusive involvement of the foveal percept in driving the updating process, and rather consider the coupling of the foveal percept with a perceptual prior in shaping the expected peripheral percept. Still, the involvement of these distinct mechanisms could depend on the feature tested, since a transfer of the updated percept at an untrained location has been found for size ^7^ but not for spatial frequency^26^.

To conclude, our study sheds light on predictive mechanisms supporting size perception stability across the visual field. A strong prior according to which objects appear smaller in the periphery contributes to the maintenance of a perceptual size difference between periphery and fovea. Such a prior should predict a specific perceived size according to eccentricity, and any deviation from this prediction would trigger the updating of the peripheral size percept. This updating should be fast^27^ to maintain stability across the visual field and ensure efficient interactions with the environment. When perception stabilized, any perceived difference could then signify a veridical change in the size of the object or of its viewing distance.

Where such a strong size prior comes from is still an open question. The perceptual bias of decreasing perceived size with eccentricity is related to the cortical structure of the visual system, with changing receptive fields from the fovea to the periphery^10^. We can only speculate that such a size prior should emerge quite early during development, based on limited cortical resolution and multimodal information, and should interact with the establishment of size constancy^28,29^. It is probably learned by long-term accumulation of stimuli statistics and is robust to changes, in order to support a stable perception by guiding the adaptive updating of peripheral percepts, as suggested by the framework of predictive adaptation^30–32^. It would be interesting to test the existence of such stabilization mechanisms for other visual features, like shape^8^ or spatial frequency^33^, that also exhibit some perceptual bias with eccentricity: would any deviating percept in the periphery be updated?

In sum, we observed the emergence of the Ebbinghaus illusion effect in the periphery. This revealed a top-down contribution to the peripheral size percept by maintaining it smaller compared to the fovea.

## Methods

### Participants

Seven participants (aged 24-44 years, including the author C.E., 3 men) took part in Experiment 1. A total of thirty-six new participants took part in Experiment2: twenty-four participants (aged 21-50 years, 6 men) in the main group (isolated and Ebbinghaus disks conditions with peripheral-foveal contingencies, see Procedure), and twelve participants (aged 20-39 years, including author CE, 3 men) in the control group (Ebbinghaus disk condition without peripheral-foveal contingencies, see Procedure). All participants had normal or corrected-to-normal vision, gave their informed written consent prior to participation, were naïve as to the purpose of the study (except C.E.) and were compensated 10 euros per hour. The national ethics committee (CPP Ile-de-France VI Pitié-Salpétrière Hospital Group) approved the protocol for the study (C13-41), in accordance with French regulations and the Declaration of Helsinki.

### Stimuli and setup

The experiments were programmed in MATLAB, using the Psychophysics and Eyelink Toolbox extensions^34–37^. Stimuli were presented in a lit room at a 52 cm-viewing distance on a 24-in monitor screen or at a 56 cm-viewing distance on a 27-in LCD monitor screen LCD (1920 ⨯ 1080 pixels, 144 Hz). Movements of the right eye were recorded using an Eyelink 1000 eye-tracker (SR Research Ltd., Mississauga, ON, Canada) operating at 1000 Hz. A chin and a forehead rest stabilized the participant’s head. Participants responded using a keyboard.

The stimuli were black disks presented in isolation or as Ebbinghaus disks on a gray background (24.8 cd/m^2^; Fig. 1a). Ebbinghaus black disks were surrounded by six white 2.1°-diameter inducers, with a 2.6° center-to-center distance. The border of each disk was blurred by a cumulative Gaussian gradient over a width of 15% of the stimulus diameter equally distributed from either border’s sides. Two vertical and two horizontal black line segments (0.35° length, 0.10° width) separated by 0.7° constituted the fixation cross.

### Experiment 1

#### Stimuli

In Experiment 1, we evaluated the apparent size of a Standard black disk, tested in three possible sizes computed by multiplying a reference surface of a 1.6°-diameter disk by a sizing factor (0.9, or 1, or 1.1; Table 1). The three corresponding Standard diameters were approximatively 1.5°, 1.6° and 1.7°. Each Standard disk was compared to a Test disk with one out of seven possible Test sizes, computed by multiplying the Standard disk surface by a sizing factor ([0.6 0.75 0.9 1 1.1 1.2 1.45], mean sizing factor=1; Table 1; refer also to Procedure). To prevent memorization of the Standard size, we also used a Lure disk that was presented instead of the Standard disk, with two possible Lure sizes computed by multiplying the Standard disk surface by a sizing factor of 0.9 or 1.1. For both Lure disks, a disk which size was equal to the Standard (sizing factor=1) was presented instead of the Test disk.

**Table 1.**
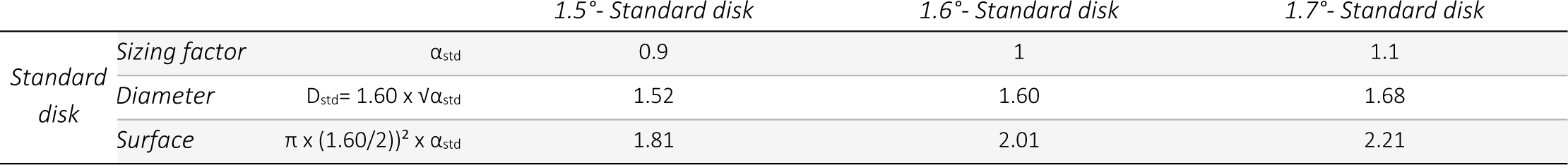
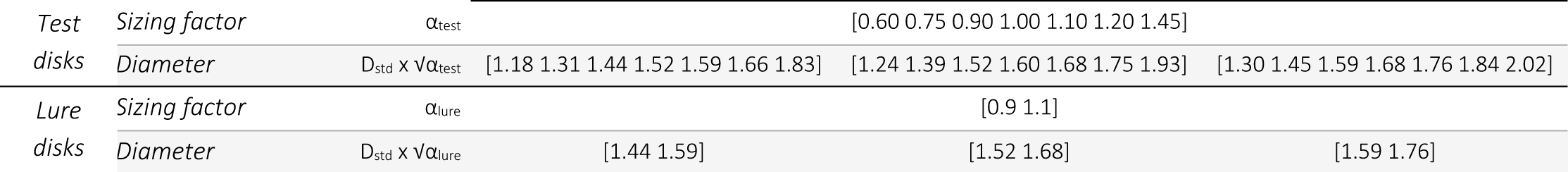
Stimuli parameters in Experiment 1. Parameters of the three Standard black disk stimuli tested in Experiment 1 (1.5°, 1.6° and 1.7°-Standard diameters). For all the Standard sizes and the corresponding Test and Lure sizes, the inducers for the Ebbinghaus disks were 2.1°-diameter white disks surrounding the central black disk at a 2.6° center-to-center distance. The three Standard disks of Experiment 1 were also used in Experiment 2 as Test disks. D stands for the diameter of the black disk, and α for the sizing factor.

#### Procedure

In Experiment 1, we used a comparison task to evaluate the apparent size of the Standard disk in two disk type conditions (isolated or Ebbinghaus disk) and at two retinal locations (peripheral or foveal) (Fig. 1a). The three Standard disks (1.5°,1.6°, and 1.7°-diameter) were tested in three separate sessions held on different days, lasting approximately 45 minutes each. In each session, the retinal location condition was manipulated in two separate blocks of 312 trials each, with a short break between them and their order counterbalanced between participants. The disk type condition was randomly interleaved within each block, with 156 trials per disk type, comprising the seven Test disks (7 Test sizes ⨯ 4 fixation positions ⨯ 5 repetitions) and the two Lure disks (2 Lure sizes ⨯ 4 fixation positions ⨯ 2 repetitions). Each trial started with a central or close to central fixation (at the center of the screen, or 1° above, or 1° below, or 1° on the left). In the peripheral condition, participants had to maintain fixation all along the trial (Fig. 1a, right panels). Fixation was controlled online and considered broken when the gaze fell outside a 1.5°-diameter region around the fixation cross; the trial was then aborted and rerun at the end of the block. After a 500-700ms fixation, a foveal black Test disk (seven possible Test sizes) replaced the central fixation cross. Simultaneously, at 12° to the right periphery appeared the Standard disk or a Lure disk, either as isolated or as an Ebbinghaus disk. Participants reported if they perceived the foveal or the peripheral black disk as largest (left or right side disk, using the left and right-arrows of a keyboard). The disks remained on the screen for 300-500ms after the participant’s response, and the next trial started 1s later. The foveal condition was identical, with the two following exceptions (Fig. 1a, left panels): the Test disk and the Standard or Lure disks appeared at 6° respectively to the left and to the right of the starting central fixation; and once the stimuli were displayed participants could freely make saccadic eye movements to bring either stimuli into foveal vision before responding.

#### Size perception data

In Experiment 1, we evaluated the apparent size of a Standard disk. Apparent size was defined as the physical size of the Test disk that should be displayed to be perceived as large as the Standard disk, and was evaluated as a ratio of the Standard size. For each Test size, we computed the proportion of trials for which the Test disk was perceived as larger than the Standard disk. Trials with Lure disks were excluded from the analysis. We fit a cumulative Gaussian function to the data (Maximum Likelihood Estimation) and determined the PSE, i.e. the apparent size of the Standard disk. Any difference in apparent size (PSE) between retinal locations or disk types would reflect a similar difference in perception: a smaller apparent size means that the standard disk was perceived as smaller.

#### Statistics

In Experiment 1, we performed a 3×2×2 repeated measures ANOVA using the open-source software JASP ^38^. Before analyzing further the interaction effect between the disk type (isolated or Ebbinghaus disk) and the retinal location (fovea or periphery) on apparent size, we conducted a Bayesian analysis ^39^ to discard the Standard diameter factor (1.5°, 1.6°, or 1.7 °-diameter) from our analysis. Bayesian analysis gave the Bayes factor (BF_10_) of model comparison between the null model and each possible model including each of the main or interaction effects. Bayes factor inclusion scores (BF_incl_) for main and interaction effects were also computed by Bayesian model averaging. To further analyze the interaction between the disk type and the retinal location, we computed the mean perceptual size ratio of the peripheral and foveal apparent sizes and compared it between the isolated and Ebbinghaus disks. We also compared the peripheral and foveal mean illusory effects, defined as the percentage of apparent size difference of the Ebbinghaus disk relative to the isolated disk, using two-tailed pair-wise and one-sample t-tests, and Bayesian analyses.

### Experiment 2

#### Stimuli

In Experiment 2, a unique 1.6°-diameter black disk served as the Reference at the fovea. In the periphery, the Ebbinghaus or the isolated black disk was displayed with one out of seven Test sizes, computed by multiplying the foveal reference surface by a sizing factor ([0.6 0.75 0.9 1 1.1 1.2 1.45], mean sizing factor=1; Test sizes computed with the sizing factors 0.9, 1, and 1.1 were identical to the three Standard sizes in Experiment 1).

#### Procedure

For the main group with peripheral-foveal contingencies, we evaluated the peripheral equivalent size of the foveal Reference with the isolated or Ebbinghaus Test disks. The experimental procedure was adapted from a study by Valsecchi and Gegenfurtner (2016) ^7^ and had three temporal phases: pre-test, middle phase, and post-test, with identical sets of trials and procedure across phases (Fig. 2a). The two disk types (isolated and Ebbinghaus disks) were tested in two separated sessions on different days, with their order counterbalanced between participants. In each session that had 357 trials in total and lasted approximately 45 min, the pre-test comprised the first 112 trials and the post-test the last 112 trials, with the seven Test disks randomly presented in the periphery (7 Test sizes x 16 repetitions). The middle phase comprised the remaining 133 trials (7 Test sizes x 19 repetitions). Each trial had two parts: first, a comparison task to evaluate the peripheral equivalent size, followed by a saccade task establishing a peripheral-foveal contingency. Importantly, during a trial, there was no change in the physical size of the stimuli. After a 500-700ms central fixation, the foveal reference disk appeared simultaneously in the center of the screen together with a peripheral Test disk at 12° to the right. The participants had to maintain fixation and answer whether they perceived the peripheral black Test disk as larger or as smaller than the foveal Reference by pressing the up- or down-arrow keyboard keys. 500-700ms after their response, the foveal reference disappeared and signaled to the participants to make an accurate saccade toward the center of the peripheral Test disk. Participants received an auditory feedback on saccade landing accuracy: a high tone indicated an accurate saccade towards the peripheral disk (see Eye Movements section for criteria) and a low tone that the accuracy might be improved. The screen was blanked 500-700ms after saccade detection (see Eye Movements section for detection criteria). On average, the peripheral Test stimulus was presented for 1725ms ± 201 (SD) prior to saccade detection, and for 566ms ± 6 (SD) at the fovea after the saccade termination (mean computed across stimulus conditions; see the Analyses section).

For the control group without the peripheral-foveal contingency, we evaluated the peripheral equivalent size for the Ebbinghaus disk in one session comprising 357 trials in total and lasting approximately 20 min. The procedure was identical, with the exception of each trial including only the first part of the comparison task, without the saccade towards the peripheral Test disk. The screen was blanked 180-270ms after the manual response.

#### Size perception data

In Experiment 2, we evaluated the peripheral equivalent size of the foveal Reference. The peripheral equivalent size was the physical size that the peripheral Test disk must have to be perceived as large as the foveal Reference disk, and was evaluated as a ratio of the Reference size. The PSE was determined by fitting a cumulative Gaussian to the data, as in Experiment 1. An increase in the peripheral equivalent size (PSE) indicated a decrease of perceived size in the periphery, since the peripheral stimulus had to be physically larger to match the foveal reference.

#### Eye movements data

In Experiment 2, several screening criteria were controlled online: (a) Central fixation was mandatory during the size comparison task and until the saccade signal (the disappearance of the foveal disk); when fixation was broken (gaze falling outside a 1.5°-diameter region around the fixation cross), the trial was aborted and rerun at the end of the phase. (b) If no saccade was made 1.5 s after the signal, the trial was also aborted and rerun at the end of the phase and a message reminded the participant to make a saccade. (c) Landing accuracy was measured within a 1°-diameter region around the saccade target and served for auditory feedback. The response saccade was detected when gaze position crossed a vertical boundary at 2° to the right of the central reference. Onset and offset of the response saccade were recorded during the trial and detected with the default EyeLink criteria defined by eye speed (30°/s) and acceleration (8,000°/s^2^). An offline analysis excluded trials according to the following criteria: (a) anticipatory answer in the comparison task (manual reaction time < 300ms, given by the time difference between the stimuli appearance and the key press time), (b) invalidated foveal view of the peripheral Test disk after the saccade towards it (gaze never focused on the peripheral disk at less than 3 degrees before its disappearance and after the saccade detection). The rates of rejection were 1.51% and 1.36% respectively in the Ebbinghaus and isolated disks conditions. Before the response saccade, the mean duration of the peripheral stimulus presentation was computed as the difference between its appearance and the saccade detection times. After the response saccade the mean foveal display duration was the time difference between the first fixation within a 3°-diameter region around the stimulus and the disappearance of the stimulus.

#### Statistics

For Experiment 2, a 2⨯2⨯2 mixed ANOVA and Bayesian analyses (Bayes factors of model comparison BF_10_ and Bayes factors inclusion scores of effects BF_incl_) were used to test the interaction effect between the disk type (isolated or Ebbinghaus disk) and the evaluation phase (pre- or post-test) on the peripheral equivalent size. We included the stimulus order factor (Ebbinghaus disk tested in the first or second session) to remove it from the following analysis. To further analyze the interaction between the stimulus and the phase conditions, we computed the updating as the percentage of peripheral equivalent size change between the pre-test and the post-test and compared it in the isolated disk condition to that of the Ebbinghaus disk condition. We also computed the percentage of peripheral equivalent size difference between the Ebbinghaus and isolated disks conditions and compared this peripheral illusory effect in the pre-test to that in the post-test, using one-sample or pair-wise t-tests and Bayesian analyses. Finally, we compared the updating in the Ebbinghaus disk condition between the main and the control group, using a mixed ANOVA.

In Experiment 2, we tested our hypothesis with two-tailed t-tests, for a decrease in peripheral perceived size of the Ebbinghaus disk between the pre- and post-test phases and for a larger updating for the Ebbinghaus disk compared to the isolated disk. We also expected in the pre-test phase of Experiment 2 to replicate the reversed illusion effect in the periphery, and to observe in the post-test the classic reduction on perceived size, with a larger effect than in the pre-test.

In both Experiments, we evaluated the normality of data distribution with the Shapiro-Wilk test when performing t-tests, and the equality of variances between groups with Levene’s test when performing mixed ANOVAs, with sphericity corrections when needed. Effect sizes were measured using Cohen’s d and η_p_^2^. We reported also the 95% confidence interval (CI) of computed mean values.

## Data Availability

The datasets generated during and analyzed during the current study are available on the Open Science Framework repository [https://osf.io/dpbu7/].

## Acknowledgement

This research received funding from the FRM grant agreement n° DEQ20150331737 to L. N., from Academy of Sciences (Neurology Lamonica Prize 2016) to L.N., and from Marie Skłodowska-Curie fellowship n° 702577 and ANR-16-CE37-0005 to T.S.M.

## Author contributions

C. E. and L. N. developed the study concept. All authors contributed to the study design. C. E. performed testing, data collection and the data analysis. All authors performed the interpretation of the results. C. E. drafted the manuscript, and T. S. M. and L. N. provided critical revisions. All authors approved the final version of the manuscript for submission.

